# Electrically controlled interaction between cable bacteria and carbon electrodes

**DOI:** 10.1101/2023.08.14.553267

**Authors:** Robin Bonné, Ian P.G. Marshall, Jesper Bjerg, Ugo Marzocchi, Jean Manca, Lars Peter Nielsen, Kartik Aiyer

## Abstract

Cable bacteria couple the oxidation of sulphide in sediments to the reduction of oxygen via long-distance electron transfer through periplasmic wires. While direct electron transfer between cable bacteria cells belonging to the same filament is a well-known phenomenon, electron transfer from the filament to electrodes has remained elusive. In this study, we demonstrate that living cable bacteria are attracted to electrodes in different bioelectrochemical systems. Carbon felt and carbon fibre electrodes poised at +200 mV against an Ag/AgCl reference attracted live cable bacteria from the sediment. When the applied potential was switched off, cable bacteria retracted from the electrode. qPCR and scanning electron microscopy corroborated this finding and revealed cable bacteria adhered onto the electrode surface. These experiments raise new possibilities to cultivate cable bacteria and utilise them for important applications in bioelectrochemical systems.

## Introduction

The ability of certain microorganisms to perform extracellular electron transfer (EET) by reducing a solid, external electron acceptor is a unique strategy for energy generation when faced with a lack of soluble electron acceptors in the immediate environment. These microbes, termed variously as electricigens, electroactive, exoelectrogens or anode-respiring bacteria use specialised mechanisms either involving redox-active outer membrane cytochromes ^1^, conductive nanowires ^2^ or soluble redox shuttles to mediate EET ^3^. Bioelectrochemical systems (BESs) exploit this unique respiratory ability of electroactive bacteria to convert chemical energy to electricity (or vice versa). Microbial oxidation of electron donor compounds provided as fuel generates electrons, which are subsequently captured by the electrode and flow in a circuit, resulting in current. BESs have provided a wealth of information about different modes of EET, especially in model electroactive bacteria like *Geobacter* ^4,5^ and *Shewanella* ^3,6^. In recent years, they have also provided useful insights into the fundamental nature of electroactivity in different classes of microbes, including gram-positive bacteria ^7,8^, pathogenic bacteria ^9–11^ and weak electricigens ^12,13^. On the applied side, BESs have branched out into several useful applications, including wastewater treatment ^14,15^, heavy metal recovery ^16,17^, biosensing of hazardous chemicals ^18,19^, powering remote sensors ^20,21^ and even monitoring and removal of viral antigens in a medical context ^22,23^. The multidisciplinary nature of this field fuses knowledge from electrochemistry, microbiology, engineering and material science, and has contributed towards important applications in diverse fields.

The discovery of cable bacteria has added a new layer of complexity to microbial electron transport processes ^25,26^. Cable bacteria are present in environments where electron donors and acceptors are spatially separated by centimetre distances ^27,28^. In order to complete metabolic reactions for energy generation, cable bacteria utilise long-distance electron transfer (LDET), in which they couple two redox half-reactions over distances far exceeding cell-length ^27,29^. Found ubiquitously in freshwater and marine sediments ^30,31^, cable bacteria oxidise sulphide in the anoxic zone and transport the electrons via conductive periplasmic proteins to the upper sediment layers to reduce oxygen or nitrate ^26,32^. A single cable bacterium filament may consist of thousands of individual cells, and may be up to 5 cm in length. A unique phenomenon associated with cable bacteria is the presence of other “flocking” bacteria around them, presumably for reducing cable bacteria via interspecies electron transfer ^33^. This warrants further investigation into whether there are mechanisms developed by cable bacteria for performing extracellular electron transfer under certain conditions.

Recently, cable bacteria were found on the anodes of a benthic microbial fuel cell, raising speculations regarding their ability to perform EET ^34^. However, further research probing the ability of cable bacteria to perform EET has been lacking. Despite the ability of cable bacteria to survive anoxic conditions without performing LDET ^35^ and the fact that periplasmic wires extracted from cable bacteria retain their conductivity in vacuum ^24^, systematic evidence for living cable bacteria to perform EET is still missing. A challenge currently faced in this aspect is that cable bacteria are not yet isolated in pure culture. In this study, we visualize and quantify cable bacteria around carbon electrodes in two different BES setups. A first conventional three-electrode cell allowed for amperometry, voltammetry, qPCR, and scanning electron microscopy (SEM) to determine the presence and impact of cable bacteria on electrodes. Another BES was specifically designed to study the interaction with electrodes in real time using non-invasive microscopy. Our data suggests that the electrode is a viable alternative electron acceptor for cable bacteria. This could help tap into the electric potential of cable bacteria to develop various applications, while also offering a chance to explore the fundamental aspects of the electron transfer process.

## Results and Discussion

### Current generation by the cable bacterium consortium

To investigate the potential electroactivity of cable bacteria, a three-electrode cell consisting of carbon felt working electrode, Ag/AgCl reference electrode and a Ti counter electrode was inoculated with freshwater sediment enriched with *Candidatus* Electronema aureum (Fig. 1a). After inoculation, the current demonstrated a sigmoidal increase (Fig. 1b), with average values ranging between 17µA and ∼78µA. This indicates the capability of the cable bacteria consortium to reduce the electrode via EET. The control cells with autoclaved sediment produced a steady background current of 4.75 **±** 0.5 µA throughout the incubation. Furthermore, the pH of the overlying water in the inoculated cells decreased over time, from 7.4 to 6.7 (supplementary table S1), while the pH of the control cells did not change. These changes arise as a result of mass transfer of protons after biological substrate oxidation, and the trend is consistent with the overall behavior of microbial three-electrode cells ^36^.

**Figure 1.**
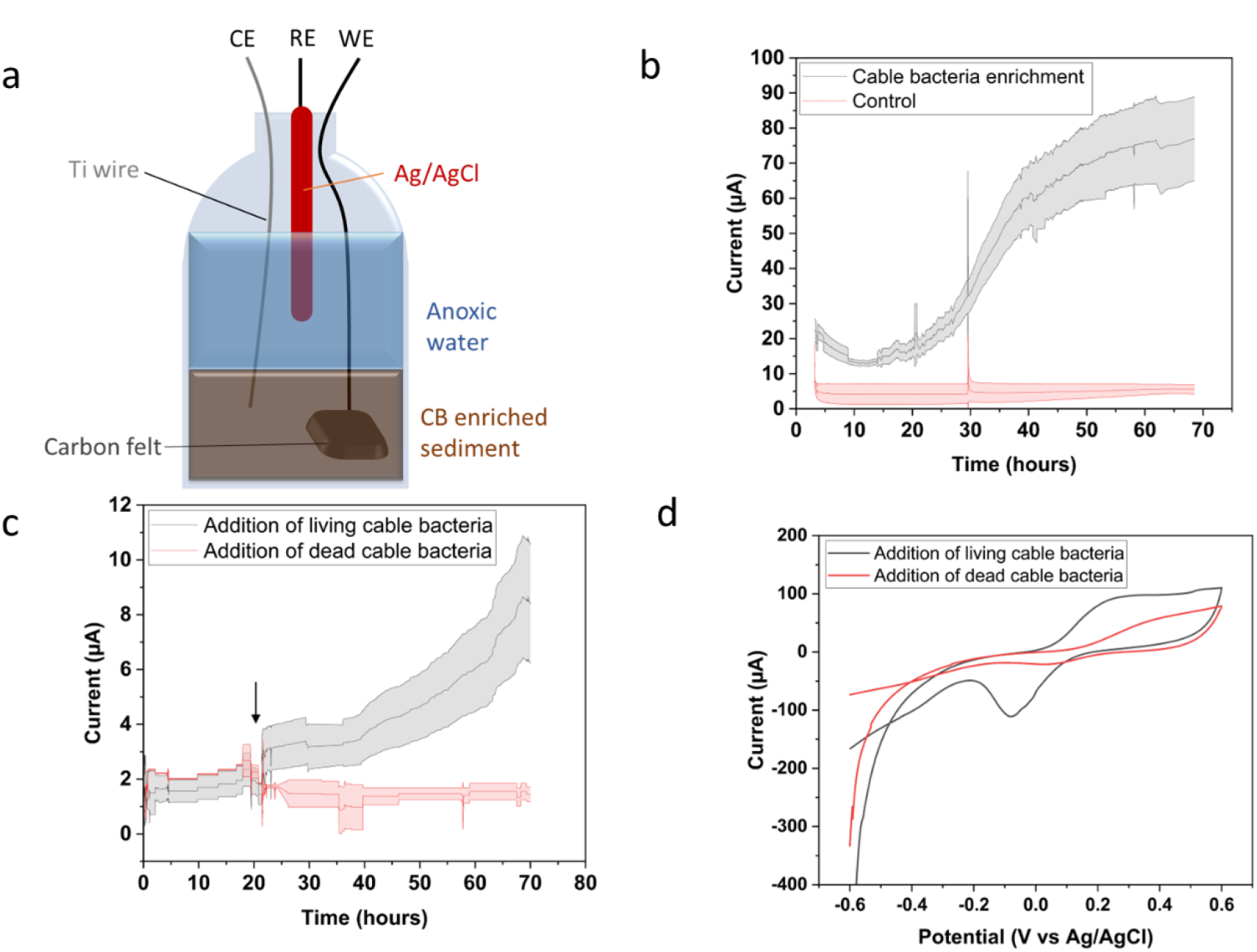
(a) Schematic representation of a sediment-based BES (b) Chronoamperometry results show a rising current within 24 hours. The data represented is the mean current value from three reactors, while error bars represent the standard deviation. (c) When isolated cable bacteria are added to autoclaved sediment (indicated by the arrow), the current rises significantly within a day (d) Cyclic voltammograms (average from three reactors) performed after experiment 1c show a redox active behaviour.

To further elucidate the specific contribution of cable bacteria to the overall measured current, another set of 3-electrode cells were inoculated with autoclaved sediment, to which approximately 10 clean cable bacteria filaments were added after fishing them from the sediment using sterilized glass hooks. Within 48 hours after addition of living cable bacteria, the current increased from ∼2µA to ∼ 8µA (figure 1c). Addition of heat-killed cable bacteria to autoclaved sediments did not increase the current. The results suggest that cable bacteria reduce the electrode in the absence of conventional electron acceptors like oxygen. However, it is essential to note that other electroactive bacteria could also have been transferred along with cable bacteria in this process.

Figure 1d demonstrates the cyclic voltammograms for the above experiment when cable bacteria are added to autoclaved sediment. Prominent redox peaks are observed at 0.22V and -0.05V, signifying the electrochemical activity of cable bacteria. While these peaks do not seem to be associated with outer membrane cytochromes typically involved in EET ^37^, they could shed light on a potential mechanism for EET.

### qPCR and SEM reveal increased presence of cables on the electrode

To assess whether the cable bacteria were abundant on the electrodes, qPCR was performed on the carbon felt working electrodes. qPCR was performed at the completion of the experiment after 72 hours, both on the carbon felt working electrodes and sediment approximately 3 cm away from the electrode. Both samples were taken in triplicates. The number of cable bacterial gene copies was significantly higher on the electrode surface as compared to the sediment (Fig. 2). Cable bacteria were on average around 100 times more abundant when a potential was applied (reactor 2) compared to the original sediment, and around 1000 times more abundant compared to the controls (C1,C2,C3) without an applied potential. With the applied potential, they were 10-1000 times more abundant around the electrode than in the bulk sediment. In the open circuit negative controls C2 and C3, a similar trend was observed, which could indicate that the unpoised carbon felt functions as an electron storage capacitor ^38^. Control C1 however showed higher abundance and deviated from the trend, which could be due to oxygen leaking into the reactor.

**Figure 2.**
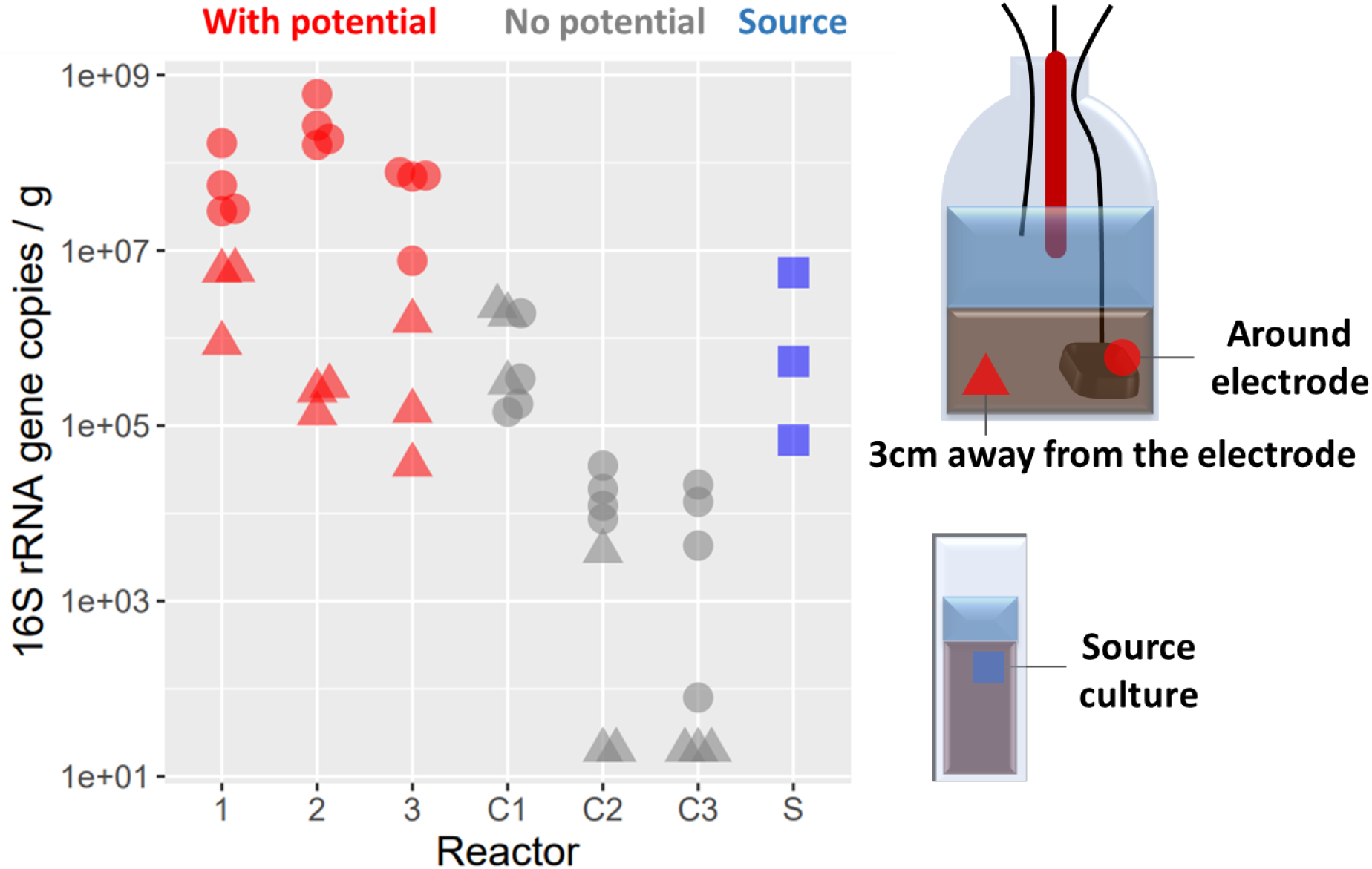
qPCR performed on the electrode reveals an increased abundance of *Ca*. Electronema aureum on the electrode surface. Reactors 1C, 2C and 3C are the controls (without applied potential), whereas 1, 2 and 3 are the experimental reactors. Source indicates the freshwater sediment containing the single strain enrichment of *Ca*. Electronema aureum

To demonstrate that the cable bacteria attached to the electrodes were living filaments, a specialized setup known as the trench slide was used. A trench slide is a microscopy slide with a central, cavity that is filled with sediment ^27^. Living cable bacteria stretch out from the sediment towards the edge to access oxygen and can be visualized via microscopy. The trench slides were filled with the sediment used in the three-electrode cell to screen the activity of cable bacteria adhered onto the carbon felt electrodes. One set of trench slides were inoculated with sediment present on the electrode surface and in the immediate vicinity of the electrodes, while the other set was inoculated with sediment 3 cm away from the electrode. Living cable bacteria filaments were observed only in the sediment collected near the electrode (Fig. S1) further substantiating the qPCR results.

To understand whether cable bacteria directly interacted with the electrode, we investigated the carbon felt working electrodes from these experiments with scanning electron microscopy (SEM) (Fig. 3). Cable bacteria were visible on all of the tested samples originating from the poised anodes (Fig. 3a-c). Some cables were coiled around the carbon felt electrode, possibly to maximize contact area with the electrode. The cable bacteria present on the electrodes had lengths exceeding 150µm. The individual cells within the cables were ∼2µm in length and cell-cell junctions were clearly observed in the SEM images, along with mineral deposits. For the negative control, where the electrodes did not have an applied potential, cable bacteria were not found on the electrode surface (Fig. 3d).

**Figure 3.**
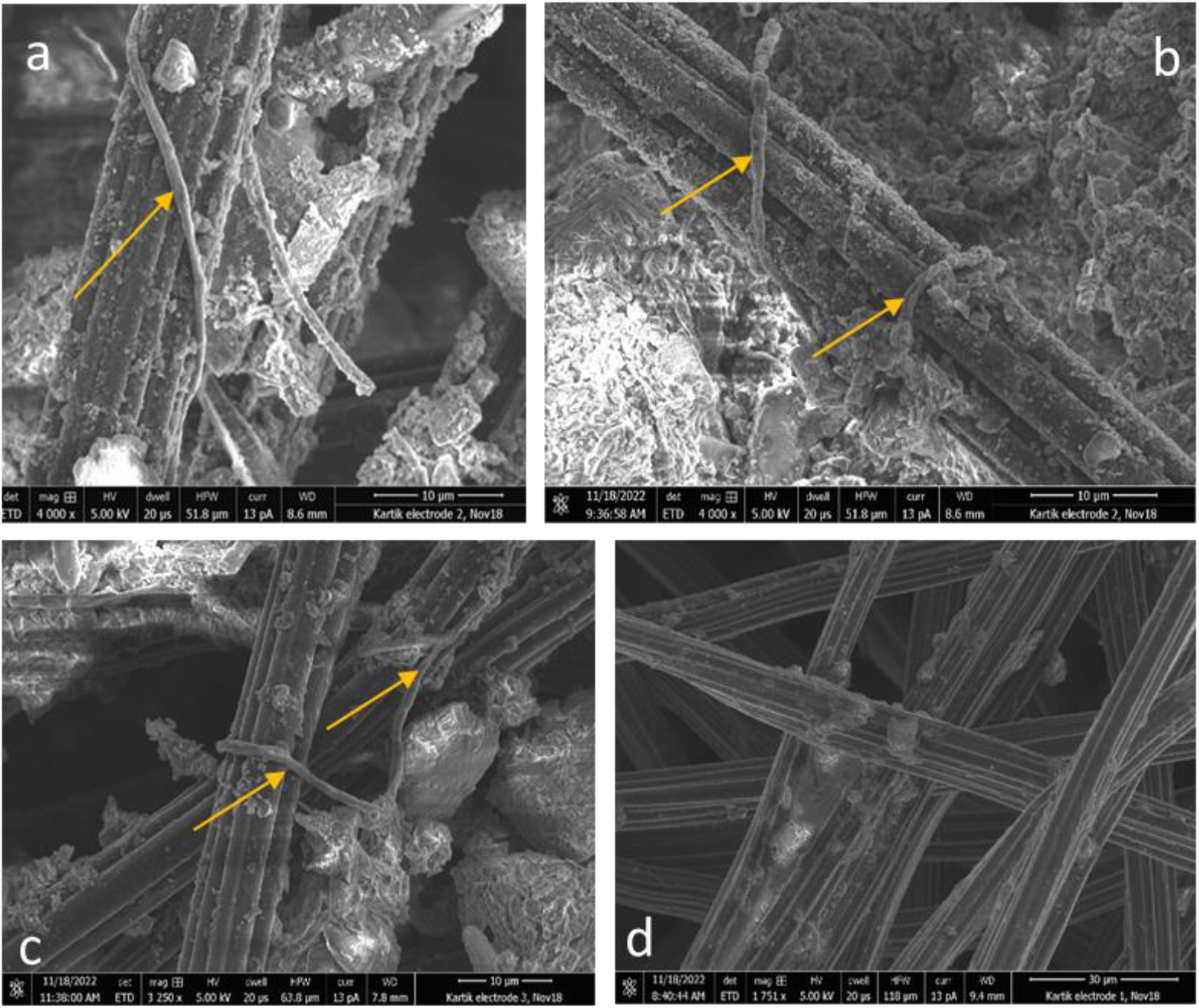
Scanning electron microscopy of the electrodes from the three-electrode cells. Images a-c show the presence of cable bacteria on the electrodes (indicated with yellow arrows), while d is the negative control (electrode without applied potential)

### Real time attachment of cable bacteria to electrodes using light microscopy

To observe the response of cable bacteria to a switchable applied potential in real time, a different configuration of the microbial electrochemical cell was constructed within a trench slide. Carbon fiber (CF) electrodes were glued parallelly on either side of the trench to function as the working and counter electrodes, and an Ag/AgCl pseudoreference electrode was introduced 3 cm from the trench (Fig. 4a). The edges of the trench slide were sealed to block oxygen diffusion in the trench slide, leaving the CF electrode as the sole electron acceptor. The bacteria started populating the poised CF electrode within a few hours after inoculation (supplementary video 1) along with simultaneous generation of current. By contrast, no cable bacteria were observed at an unpoised CF electrode (Fig. 4b). While most of the cable bacteria aggregated as white filamentous mass around the electrode, some retained their connection to the sulfide rich sediment, to actively couple their electron donor and acceptor (see supplementary video 2 and video 3 to compare the poised and unpoised carbon fibers). It is however possible that sulfide diffused out of the sediment, giving the aggregated cable bacteria access to an electron donor.

**Figure 4.**
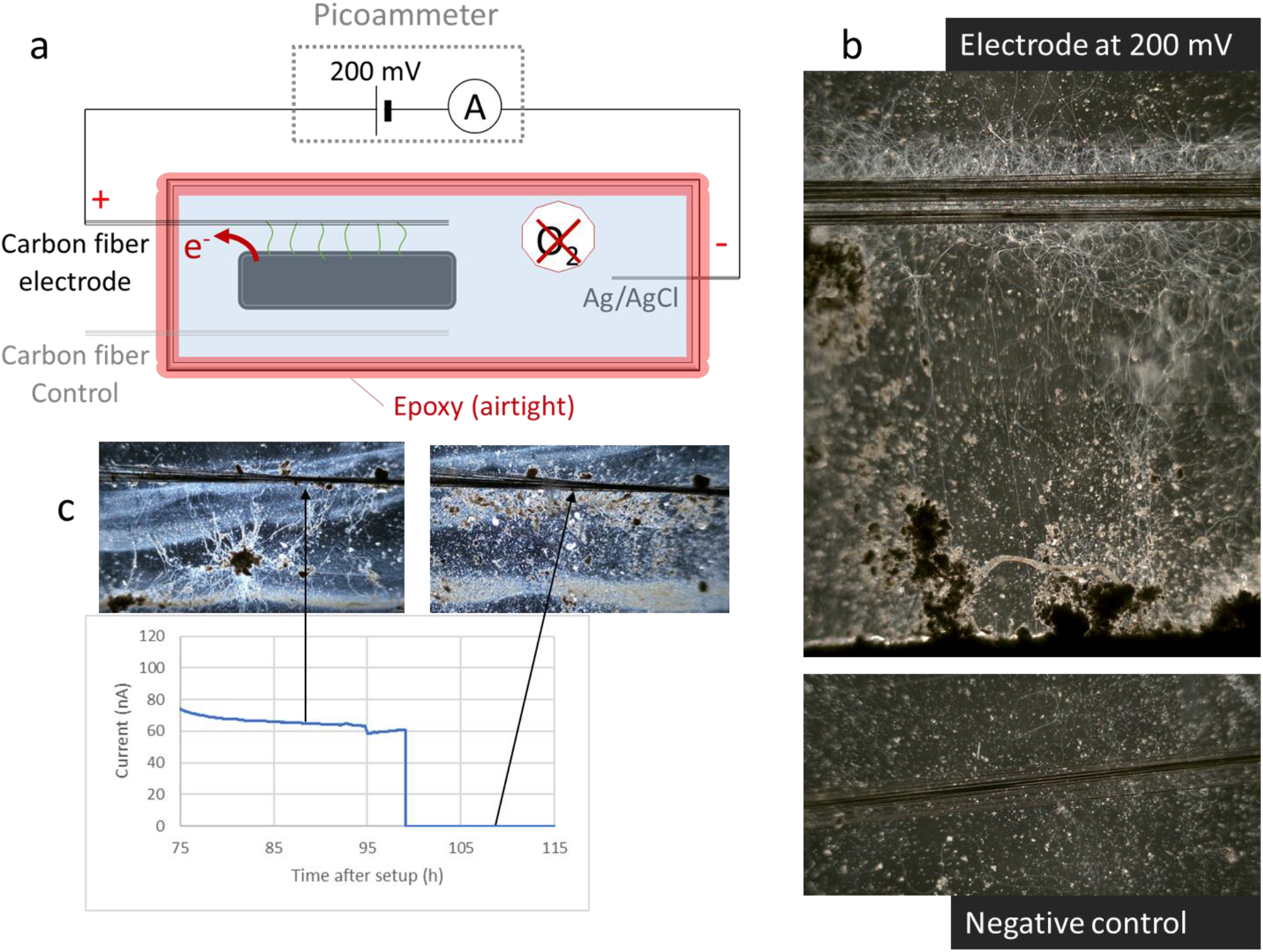
(a) An electrochemical cell on a microscopy slide makes it possible to study real time interactions with a carbon fiber electrode. (b) Cable bacteria populate the poised carbon electrode as a white filamentous mass, with some of them crossing between the sediment and the electrode. The unpoised electrode did not contain cable bacteria. (c) When the potential is switched off at the carbon electrode, cable bacteria retracted from the electrode surface.

In a different trench slide reactor of the same configuration, the current was monitored over 100 hours of operation, and showed a slowly increasing current that reached values between 50 and 100 nA (data not shown). The poised electrode contained several cable bacteria at the electrode stretching from the sediment (Fig. 4c). After 98h of operation, the applied potential was switched off, and cable bacteria retracted from the electrode in a matter of 10 hours. (Fig. 4c). This indicates that cable bacteria stay alive in an anoxic environment for days in the presence of a solid electron acceptor. Similarly, when the unpoised electrode was poised after 5 days of incubation, cable bacteria reappeared on the electrode after approximately 2 hours (supplementary figure S2), indicating their ability to survive the absence of suitable electron acceptors.

The open question remains whether cable bacteria make direct contact with the electrode or reduce the electrode through mediators. First indications of a direct connection can be seen through a timelapse of cable bacteria interacting with electrodes, where they seem to pull on the carbon fibers over time (supplementary video 4). Additionally, when the microscopy slide was shaken during operation, the bacteria retained their connection with the carbon fibers, indicating robustness of their connection to the electrode (supplementary video 5). Although these results point towards a physical connection between cable bacteria and electrodes, this does not necessarily indicate the presence of EET pathways in cable bacteria.

### Potential Mechanism of EET

The attraction of cable bacteria towards electrodes only in the presence of an applied potential lends credence to a putative EET mechanism by cable bacteria. The mechanism of potential EET to the electrode remains an open question, despite supporting evidence for direct and mediated electron transfer. On one hand, the cyclic voltammetry results indicate the presence of redox mediators. On the other hand, SEM and microscopy results of cable bacteria coiled around the electrode, together with previous results from Reimers et al ^34^ seem to point towards a direct electron transfer. Several cytochromes recently identified in cable bacteria ^27,39^ could also potentially be implicated in extracellular electron transfer, though their structural and functional role remains to be studied.

It is important to note that since cable bacteria have not yet been isolated in pure culture, the single-strain enrichments in sediments therefore still constitute a “mixed” community. Several of the bacteria flocking around cable bacteria ^33^ were found to belong to *ɣ-proteobacteria* (e.g. *Sideroxydans, Rhodoferax*), which have been previously confirmed to be electroactive ^40,41^. While some of the current generated is certainly due to these bacteria, the presence cable bacteria on electrodes and the qPCR results provide reasonable evidence of putative EET by cable bacteria. It also remains to be seen whether these flocking bacteria could perform interspecies electron transfers with cable bacteria for subsequently reducing the electrode.

In the natural setting, the main physiological relevance of EET could be to help cable bacteria to survive conditions where oxygen and nitrate are either depleted or unavailable. Iron and manganese-based minerals are amply present in the sediment, which could potentially be used as terminal electron acceptors under oxygen/nitrate-limiting conditions. *Vibrio natriegens*, recognized to be among the fastest respiring microbes due to its rapid oxygen consumption, has also evolved a strategy to perform EET under low oxygen conditions ^42^. Such a hybrid model could similarly benefit cable bacteria and aid in their survival under anaerobic conditions.

Overall, these findings indicate that cable bacteria, in the absence of a soluble electron acceptor, tend to migrate towards an electrode to reduce it, thereby generating electric current. However, it is not yet clear as to whether cable bacteria can actively divide and grow in such an electrogenic mode over long periods of time, though it has previously been suggested that cable bacteria can grow in anoxic conditions ^43^.

## Conclusions

Cable bacteria were found to be attracted to carbon electrodes and remain active in response to an applied potential in different bioelectrochemical systems. This work paves the road for similar bioelectrochemical experiments to elucidate the magnitude and mechanism of electron transfer from cable bacteria to the electrode as well as the behavioral triggers and any associated growth. Successful growth of cable bacteria in bioelectrochemical systems will enable the development of useful applications like bioremediation of soil contaminants, carbon capture and the subsequent production of chemicals.

## Materials and Methods

### Construction of the three-electrode cell

Three-electrode cells were constructed from glass serum bottles (70 ml volume). The inoculum consisted of single strain enrichment of *Ca*. Electronema aureum in freshwater sediment as described by Thorup et al ^44^, with other associated bacteria present in the sediment. 10g of sediment was weighed and added into the glass bottle, followed by addition of N_2_-sparged lake water. For the controls, this sediment was autoclaved. The working electrode consisted of carbon felt (dimensions 2 cm x 2 cm x 0.3 cm), a commercial Ag/AgCl electrode (REF201 Radiometer Analytical, Denmark) constituted the reference electrode, and a titanium wire was used as the counter electrode. The working electrode was buried 2 cm in the sediment, whereas the reference and counter electrodes were positioned in the overlying water (Fig. 1a). The electrodes were connected to a potentiostat (MultiSens 4, PalmSens, Netherlands), and a potential of 200 mV was applied to the working electrode (Fig. 1b). In an additional set of experiments (Fig. 1c), a similar 3-electrode setup was prepared with autoclaved sediment. Around 10-20 cable bacteria were extracted with glass hooks from natural freshwater sediment, cleaned in milli-Q water and added to cell. Cyclic voltammetry was performed before and after cable bacteria addition. The potential window was -0.6V to +0.6V at a scan rate of 10 mV s^-1^. Data was recorded using the MultiTrace software.

### qPCR

For qPCR, primers specific to the *Ca*. Electronema aureum GS strain were used (GS68F: 5’-GGTAGTTTCCTTCGGGGGAC-3’, GS178R: 5’-CTTCGGCAATGCGGCGTATC-3’), which were designed based on the 16S rRNA gene for strain from its genome using the Arb PROBE DESIGN tool ^45^ with the Silva Ref NR 99 database version 132 ^46^. DNA was extracted from sediment and electrode material (0.5 g per sample) using the DNeasy PowerLyzer PowerSoil Kit (Qiagen) following the manufacturers’ instructions. Quantification of cable bacteria was carried out using quantitative PCR with 2µL of DNA extract added to each 20 µL reaction along with RealQ Plus 2x Master Mix Green (Ampliqon, Odense, Denmark), 0.1% BSA, 0.5 pmol/µL of each primer. A synthetic standard containing base positions 1-200 from the 16S rRNA gene sequence was used for the standard curve (GenScript, Rijswijk, Netherlands). The qPCR was carried out on a Strategene Mx3005P qPCR machine (Agilent Technologies, Santa Clara, CA, USA) with an initial 15-minute activation step at 95ºC, followed by 45 cycles with 30 s at 95ºC, 30 s annealing at 60ºC, 20 s elongation at 72ºC, and 15 s acquisition at 95ºC. A melting curve from 60ºC to 95ºC confirmed the absence of secondary amplicons.

### Scanning Electron Microscopy (SEM)

After conclusion of the experiment, the carbon felt working electrode was carefully removed and SEM was performed on the electrode. The instrument consisted of a dual beam FIB/SEM 2D Versa (Thermo Fisher). Acquisition parameters included a high vacuum of 5 kV, a working distance of 10 mm and a current of 13 pA.

### Construction of the trench slide three-electrode cell

In addition to the above setup, a three-electrode cell was constructed within a trench slide system (Fig. 4a). The trench slide was a glass slide containing a central, 1 mm deep, rectangular cavity for inoculating sediment ^27^. Two bundles of carbon fiber (CF) wires (10 µm diameter, T300SC, Toray, Japan) were glued parallelly on each side of the trench at a distance of 5mm. The CF wire on one side functioned as the working electrode, while the one on the other side was left unpoised as a negative control. At a distance of 20mm from the trench, a chlorinated Ag wire was introduced to function as the pseudo-reference and counter electrode. The central trench was filled with sediment enriched with *Ca*. Electronema aureum GS, followed by addition of N_2_-sparged lake water over the sediment. A coverslip was carefully placed on top and the trench slide was incubated at 25℃ for 4 hours to enable cable bacteria to come out from the sediment. After incubation, epoxy (Loctite instant mix, Henkel, USA) was applied around the edges of the cover slip to block oxygen penetration into the trench. Using a picoammeter (Unisense, Denmark), a potential of 200mV was applied between the working electrode and the Ag wire while simultaneously measuring the current. Data was logged using the Unisense software SensorTrace.

### Microscopy of the CF electrode

The trench slide three-electrode cell was investigated via confocal microscopy (Zeiss Observer Z1, Zeiss) to study the attachment of cable bacteria onto the electrodes.

## Acknowledgements

We thank Jesper Wulff, Ronny M. Baaske and Lars Borregaard Pedersen for their technical assistance with qPCR and trench slide experiments, and Pia Bomholt Jensen for operation of the scanning electron microscope. This work was supported by Danish National Research Foundation (Center for Electromicrobiology, DNRF136; LPN), Villum Fonden (IM) and the European Union’s Horizon research and innovation program under the Marie Sklodowska-Curie grant agreement (project 101109777; KA).

## Author Contributions

All authors conceived the study, KA and RB performed the experiments and wrote the first draft. IPGM helped in analysis of the data. All authors contributed to the final version of the manuscript.

