## Supplementary Information for "Electrically controlled interaction between cable bacteria and carbon electrodes"

| Time (hours) | pH change in the overlying water |
| --- | --- |
| 0 | 7.4 |
| 24 | 7.1 |
| 48 | 6.9 |
| 72 | 6.7 |

Table S1: pH changes in the overlying water of the three-electrode cell. The pH reduced due to the electrochemical activity of the microbes, leading to mass transfer of protons.

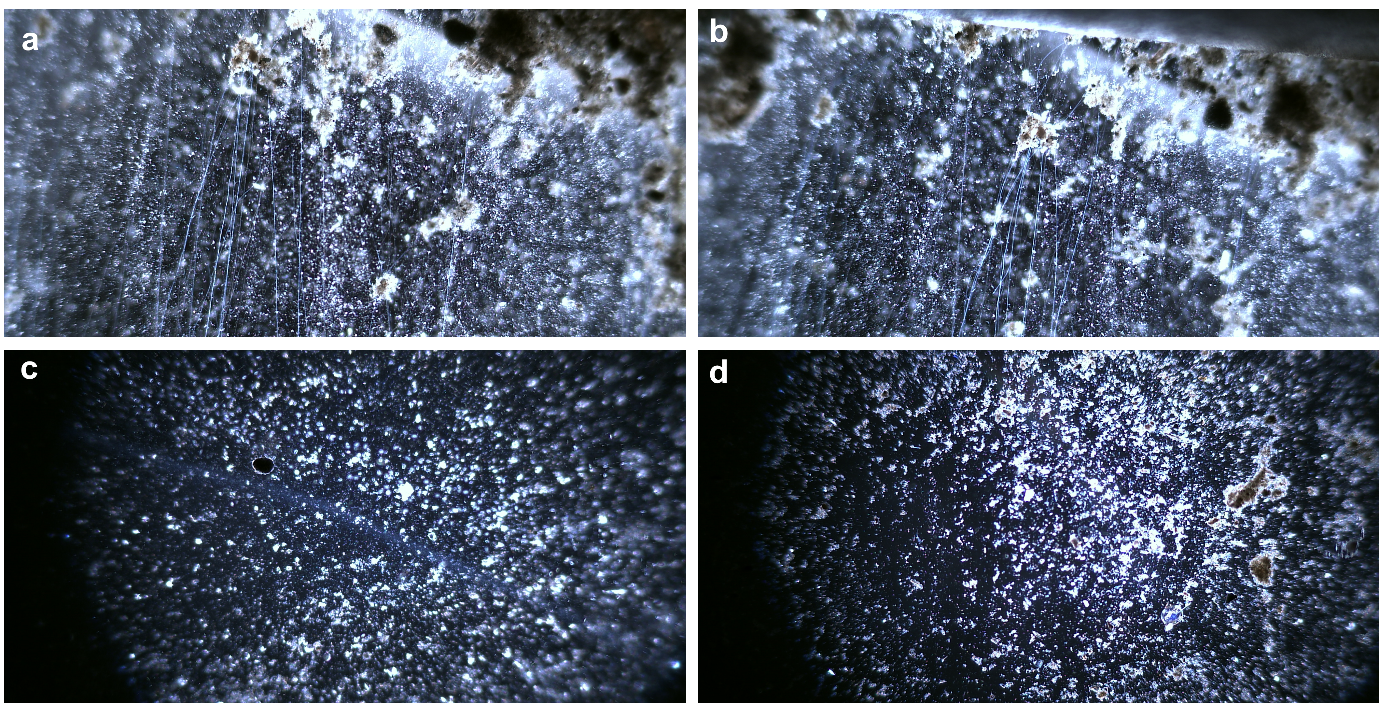

Figure S1: Migration of cable bacteria towards the electrode. After the classical BES experiment, the sediment on the electrode and away from the electrode was used to prepare trench slides. Cable bacteria were present in the sediment at the electrode (a, b), while no cable bacteria were detected in sediment away from the electrode (c, d).

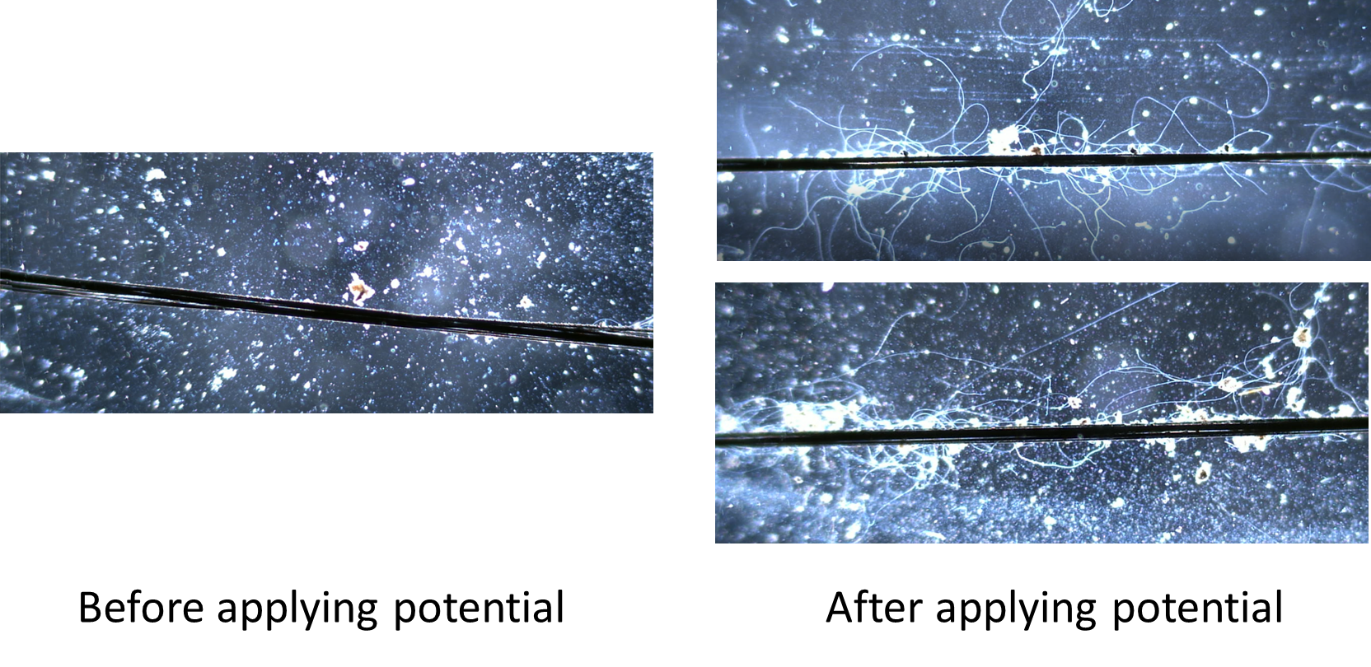

Figure S2: The carbon fibre electrode before and after applying a potential. The poised electrode attracts cable bacteria.

Video links

Video 1: [Cable bacteria populating the poised electrode](https://www.dropbox.com/s/r8v4kt0alwir7i7/VideoS1_CEBBESpaper_AiyerBonn%C3%A9_PopulatingPoised3.mp4?dl=0)

Video 2: [Cable bacteria at poised electrode](https://www.dropbox.com/s/zdrc0c8ynsdb40u/Poised%20electrode.avi?dl=0)

Video 3: [Unpopulated unpoised electrode](https://www.dropbox.com/s/ds8f7w2z631jr3m/Unpoised%20electrode.avi?dl=0)

Video 4: [Cable bacteria pulling at carbon fiber](https://www.dropbox.com/s/hjxvoqin063dd6m/VideoS4_CEBBESpaper_AiyerBonn%C3%A9_PullingCF2.mp4?dl=0)

Video 5: [Robustness of cable bacteria attachment to electrode](https://www.dropbox.com/s/ejt0p35tyiynhsp/CB%20Electrode%20Shaking.avi?dl=0)

Upon applying a potential, cable bacteria start populating the electrode in a matter of hours (video 1). The poised electrode attracts significantly larger numbers of cable bacteria, which are present on the electrode (video 2 and 3). In the unpoised electrode, this phenomenon is not observed. In video 4 and 5 we demonstrate indication of direct attachment of cable bacteria to the electrode.
